# DeepRF: Ultrafast population receptive field mapping with deep learning

**DOI:** 10.1101/732990

**Authors:** Jordy Thielen, Umut Güçlü, Yagmur Güçlütürk, Luca Ambrogioni, Sander E. Bosch, Marcel A. J. van Gerven

## Abstract

Population receptive field (pRF) mapping is an important asset for cognitive neuroscience. The pRF model is used for estimating retinotopy, defining functional localizers and to study a vast amount of cognitive tasks. In a classic pRF, the cartesian location and receptive field size are modeled as a 2D Gaussian kernel in visual space and are estimated by optimizing the fit between observed responses and predicted responses. In the standard framework this is achieved using an iterative gradient descent algorithm. This optimization is time consuming because the number of pRFs to fit (e.g., fMRI voxels) is typically large. This computation time increases further with the complexity of the pRF model (e.g., adding HRF parameters, surround suppression and uncertainty measures). Here, we introduce DeepRF, which uses deep convolutional neural networks to estimate pRFs. We compare the performance of DeepRF with that of the conventional method using a synthetic dataset for which the ground truth is known and an empirical dataset. We show that DeepRF achieves state-of-the-art performance while being more than 3 orders of magnitude faster than the conventional method. This enables easier and faster modeling of more complex pRF models, resolving an important limitation of the conventional approach.

## 1 Introduction

In visual neuroscience the investigation of receptive field properties has been one of the major advancements in understanding the visual system. Receptive field properties (i.e., location and size of the receptive field) provide a way to construct the preferred positions in the visual field of each recording site, from which visual field maps can be constructed. Furthermore, by creating a model for the receptive field one can predict neural responses to different stimuli. Recording techniques that are used in investigating human receptive field properties often take the responses of many neurons.

In creating receptive field models at this scale, one refers to the model as a population receptive field (pRF) model as it measures the pooled response from several million neurons [29].

In humans, functional magnetic resonance imaging (FMRI) provides an excellent instrument to measure brain activity as a response to visual stimuli and is often used in visual field mapping. The conventional pRF mapping approach proceeds by specifying a parameterized model (and estimating the parameters per voxel using gradient descent [30]. Often, this receptive field model is taken to be a symmetric two-dimensional Gaussian [31]. A wide variety of extensions to this basic approach has been developed such as incorporating surround suppression with a difference of Gaussians (DoG) model [36], a method without any explicit assumption on the shape of the receptive field [18], compressive spatial summation [13], Bayesian extensions of the conventional method [21, 35] and connective field modeling [7].

The pRF approach has been used to map visual cortex using retinotopy, specifically to draw regions of interest (ROIs) like dorsal and ventral V1, V2 and V3 [25], to map color throughout visual cortex [32], to map numerosity in parietal areas [10, 9], to map auditory properties throughout temporal cortex [27] and to investigate a wide range of phenomena like scotomas [6], attention [28, 16], shape perception [17], autism spectrum disorder [23], Alzheimer’s disease [2] and blindness [19].

A major bottleneck with conventional pRF mapping is the time required to compute a pRF map, which easily takes multiple hours per subject. This not only wastes computational resources and valuable time, but also precludes the estimation of more complex receptive field models that have more free parameters.

We aim to improve the methods to fit pRF models to allow faster whole-brain model fitting. We show that our proposed method is indeed faster than the baseline method while remaining on par in terms of performance. The proposed method enables faster model fitting as well as experimentation with and comparison between more complex models.

## 2 Methods

### 2.1 Forward model

We use a forward model that characterizes BOLD activity over time of an arbitrary MRI voxel. The forward model generates a voxel time series *y*(*t*) at time *t* as the sum of the signal of interest *p*(*t*) and the additive noise *ξ*(*t*):

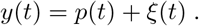

The forward model was implemented in Python (version 3.7.3).

#### 2.1.1 Signal component

The signal component is defined by two sources: the convolution of the heamodynamic response function (HRF) *h*(*t*) with the population response *r*(*t*):

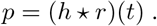

We use the double gamma HRF defined as:

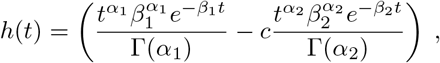

with *α*_1_ = 5 + *δ*, *α*_2_ = 15 + *δ*, *β*1 = *β*_2_ = 1 and *c* = 0.1. Here, *α*_1_ and *α*_2_ represent the time of the peak and undershoot of the HRF with a delay *δ*. The population response *r*(*t*) is the response between the stimulus *s*(*x, y, t*) and the static population receptive field *g*(*x, y*):

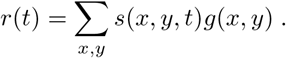

The stimulus function *s*(*x, y, t*) is a binary indicator function and denotes the presence of the stimulus at location (*x, y*) at time *t*. The population receptive field is modelled as a 2D isometric Gaussian:

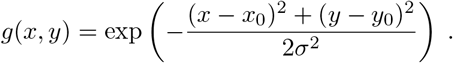

The goal is to estimate the receptive field location (*x*_0_, *y*_0_), receptive field size *σ* and HRF delay *δ*.

#### 2.1.2 Noise component

The noise component *ξ*(*t*) is based on other simulation packages (e.g., NeuRosim [33] or SimTB [5]). Hyperparameter settings are taken from [34]. For an extensive review on MRI noise sources see [20]. The noise component *ξ*(*t*) contains five different sources: low frequency noise, physiological noise, system noise, task-related noise, and temporal noise, with an amplitude of *ν*.

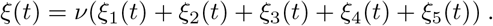

Low frequency noise models the drifts in the signal that are caused by slow fluctuations in the scanner hardware. This noise source is modeled on the basis of discrete cosine waves *θ* from the TR up to 128 seconds:

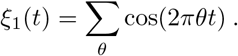

The physiological noise comprises cardiac and respiratory artefacts, which are typically at 1.17 Hz and 0.2 Hz respectively:

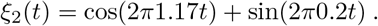

The system noise models the measurement noise, which is typically modeled as white noise:

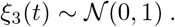

The task-related noise adds noise relative to neural activity evoked by the task as well as residual noise from head motion. It is also modeled as white noise, but only at those time points where the task is ‘active’:

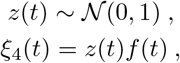

where *f* (*t*) is an indicator function that is 1 when the task is active at time *t* and 0 otherwise. Finally, the temporal noise contains the auto correlation of the signal as we are dealing with repeated measures. This is modeled as a first-order autoregressive process:

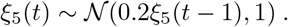

### 2.2 Conventional approach

The baseline method is the coarse-to-fine method originally proposed by [4]. Under this framework, the observed response *y*(*t*) at time *t* is modeled as:

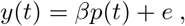

where *p*(*t*) is the predicted response, *β* is a scaling factor to account for the unknown units of the observed response and *e* is the measurement noise. The predicted response is the convolution of the heamodynamic response function (HRF) *h*(*t*) and the modeled population response *r*(*t*):

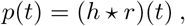

where *h*(*t*) is modeled as the double-gamma response function as defined above with delay *δ*. The modeled population response *r*(*t*) is the combination of the static receptive field properties of the population *g*(*x, y*) and the stimulus *s*(*x, y, t*) at time *t*:

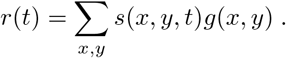

The *s*(*x, y, t*) is an indicator function that is 1 when there is a stimulus at coordinates (*x, y*) at time *t* and 0 otherwise. Here, *g*(*x, y*) is modeled as a 2D isotropic Gaussian:

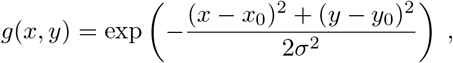

where *x*_0_, *y*_0_ and *σ* are the parameters of the pRF model describing its receptive field location (*x*_0_, *y*_0_) and receptive field size *σ*. Then, the quality of the predicted response can be measured as the sum of squared errors:

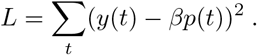

The optimal parameters *x*_0_, *y*_0_, *σ* and *δ* are estimated by a coarse-to-fine search that minimizes *L*. In the coarse stage, a grid search is performed with several combinations of plausible parameters. The combination of parameters with the smallest *L* in the coarse fit is used as the initialization for the fine stage. In the fine stage, the optimization fine-tunes the parameters further using a gradient descent approach. In the current study we used the Popeye library, which is a Python implementation of the baseline method [3]. Note, that Popeye also fits *β*, which should be 1 as we normalize the observed response *y*(*t*) before subjecting it to any model. Note that the computationally costly coarse-to-fine approach needs to be performed for every voxel.

### 2.3 DeepRF approach

Our proposed method employs deep learning to optimize the parameters of the pRF, therefore conveniently called DeepRF. DeepRF makes use of a deep convolutional neural network (CNN) that receives the observed response as input and predicts the parameters of the pRF so that the fit between observed parameters and predicted parameters is optimized. Theoretically, this can be interpreted as a forward amortized inference procedure [1].

As an architecture, we use the 50-layer preactivation residual neural network (ResNet) [11]. Since we are dealing with 1D time series as inputs instead of 2D images, we used 1D convolutions instead of the original 2D convolutions. The ResNet was implemented in Python (version 3.7.3) using the Chainer library (version 6.0.0) for artificial neural networks.

As an objective function, the network minimized the mean squared error (MSE) loss function between predicted pRF parameters (*x*_0_, *y*_0_, *σ*) and HRF parameter (*δ*) and the ground truth.

For optimization, we used Adam with hyperparameters *α* = 0.0001, *β*_1_ = 0.9 and *β*_2_ = 0.999 [15]. The ResNet was trained in 500 epochs, each of which presented 1024 newly drawn samples from the synthetic data generator in batches of 128 samples to the model (i.e., the model saw a total of 512.000 samples).

Note that, in contrast to the conventional approach, we only need to run gradient descent once during model training. That is, DeepRF model parameters are estimated uisng simulated fMRI timeseries generated under randomly chosen pRF parameters. Subsequently, the trained model can be used to recover pRF parameters of any voxel using a rapid forward pass. The only requirement is that the experimental design with which the simulated data were generated to train the model is the same as the design that was used to collect the empirical data.

### 2.4 Experimental design

The performance of the DeepRF approach was compared with that of the conventional approach using both simulated and empirical data. To this end, we used a classical pRF mapping protocol.

Stimuli were expanding and contracting rings and clockwise and counter-clockwise wedges displayed on a mean-luminance gray background (see Figure 1). Both rings and wedges were filled with a radial colored checkerboard texture and changed colors to its complementary luminance at 5 Hz.

**Figure 1:**
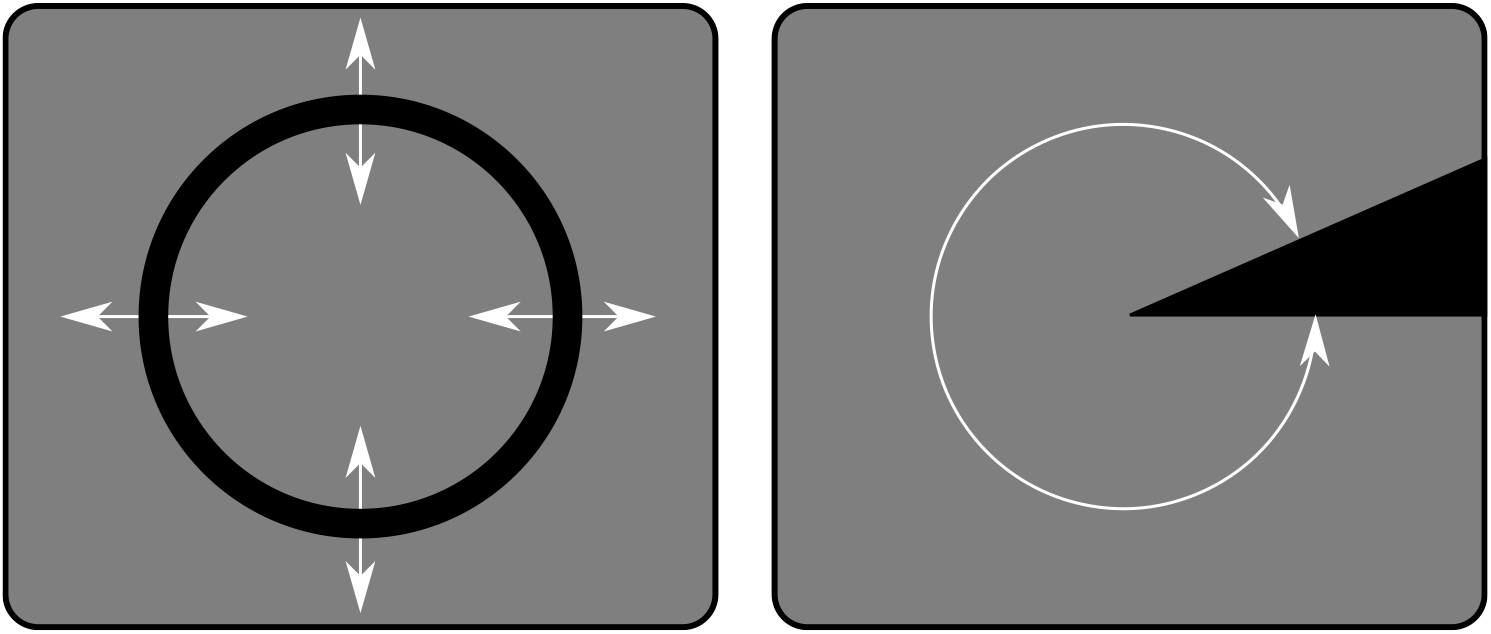
Stimuli were expanding and contracting rings (left panel) and clockwise and counter-clockwise rotating wedges (right panel). Both stimuli in both directions traveled in 16 uniform steps over the entire screen. Image adapted from [24].

The four stimuli were presented in four isolated runs. Each run lasted 180 s, starting with 4 s and ending with 16 s of pure fixation periods (i.e., no stimulus). In between, five stimulus cycles of 32 s were presented. The expanding and contracting ring had a width of 0.95 visual degree and were not scaled with the cortical magnification factor. The clockwise and counter-clockwise moving wedge had an angle of 22.5 degrees. All stimuli traveled across visual field in 16 equal steps of 2 s, making up cycles of 32 s. Each of five cycles repeated the same procedure for the specific stimulus.

This protocol was used to train the DeepRF model as well as to collect empirical data.

### 2.5 Empirical data

For empirical validation, we used the Study Forrest dataset [8]; specifically its extension to retinotopic mapping [24].

#### 2.5.1 Participants

Retinotopic mapping data was acquired for 15 right-handed participants (age 21-39 years, 6 female). For each of the participants, normal visual function was verified by means of individual eye visual acuity and visual field sensitivity tests. In this study, we used Participant 2 only.

#### 2.5.2 Experimental setup

Stimuli were presented on a 26.5 by 21.2 cm rear-projection screen inside the bore of the magnet using a LCD projector at a resolution of 1280 by 1024 pixels at a frame rate of 60 Hz. Participants perceived the stimuli via a mirror mounted on the head coil at a viewing distance of 63 cm. Stimuli were presented using PsychoPy version 1.79. During the runs, participants maintained fixation at a circle with a radius of 0.4 degrees of visual angle. To keep participants attentive, individual characters of 0.5 visual degree were presented at 1.5 Hz in the fixation circle. These characters were excerpts from song lyrics which participants had to read. After a run, participants were asked two-option forced choice questions about these lyrics to verify attention.

#### 2.5.3 MRI acquisition

Anatomical scans were T1-weighted images with 274 sagittal slices (field-of-view (FoV) of 191.8 by 256 by 256 mm) with voxel size of 0.7 mm with an 384 by 384 in-plane reconstruction matrix (0.67 isotropic resolution) recorded with a 3D turbo field echo (TFE) sequence (repetition time (TR) of 2.5 seconds, inversion time (TI) of 900 ms, flip angle of 8 degrees, echo time (TE) of 5.7 ms, bandwidth of 144.4 Hz/pix, Sense reduction AP 1.2, RL 2.0) [8].

Functional scans were T2*-weighted echo-plannar images (gradient-echo, repetition time (TR) of 2 s, echo time (TE) of 30 ms, flip angle of 90 degree, bandwidth of 1943 Hz/pix, parallel acquisition with sensitivity encoding (SENSE) reduction factor 2 acquired with a 3 Tesla Philips Achieva dStream MRI scanner and a 32 channel head-coil [24]. Whole-brain volumes were recorded in 35 axial slices (thickness 3.0 mm) with 80 by 80 voxels (3.0 by 3.0 mm) of in-plane resolution, 240 mm field-of-view (FoV), anterior to posterior phase encoding direction, 10% inter-slice gap, recorded in ascending order.

#### 2.5.4 Data preprocessing

Functional scans were motion corrected and coregistered to the anatomical scan using the SPM toolbox. Individual runs were further detrended, spectrally filtered and converted to percent of signal change by division of the pre-detrended mean. Finally, we extracted a visual ROI as defined by the Study Forrest dataset.

### 2.6 Analyses

Model performance was analysed as follows. First, we trained the DeepRF model using synthetic data generated under the described experimental protocol. Next, the conventional and DeepRF approaches were evaluated using another set of synthetic data. Model performance was compared by comparing predicted parameters with ground truth parameters. Finally, parameters estimated from empirical data were compared with each other. In addition to the DeepRF and Popeye models, we also tested a combined version, where DeepRF was used as the coarse estimation to be fine-tuned by Popeye.

## 3 Results

### 3.1 Model estimation

Training of the DeepRF model was performed on 512.000 unique samples. Training was done over 500 epochs each presenting 1024 new samples in batches of 128 samples. Training was performed on a GPU and took 3.1 hours to complete. Training loss consistently decreased over epochs (Figure 2).

**Figure 2:**
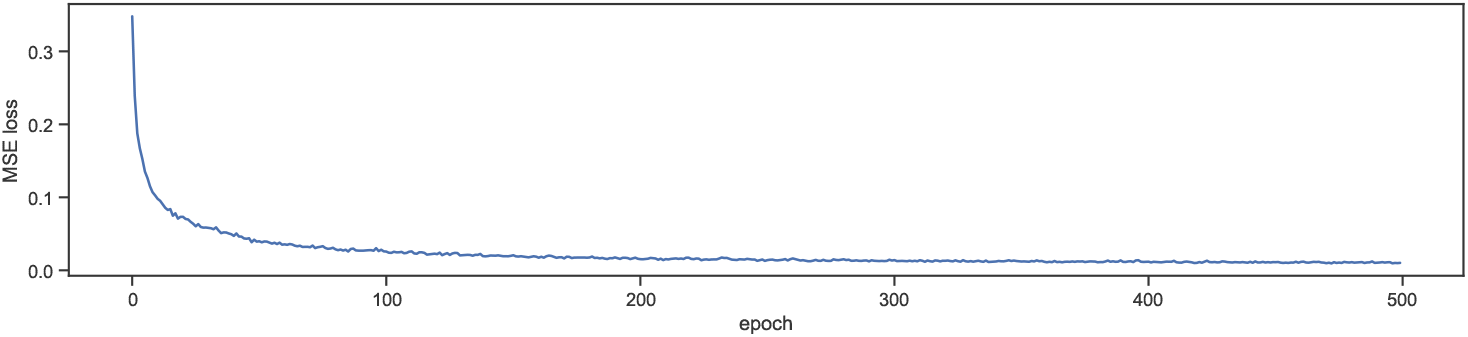
Mean squared error loss of DeepRF over training epochs.

We validated DeepRF and Popeye on newly generated syntetic data as well as empirical data. The results are outlined below. In addition to the two frameworks, we ran DeepRF as the coarse estimation to be fine-tuned by Popeye. In terms of computational speed, DeepRF was 1868 times faster than Popeye on the synthetic dataset and even 6178 times faster on the empirical dataset. Using DeepRF as initialization for Popeye resulted in a factor of two faster estimations (Figure 3). Note, both DeepRF as well as Popeye were ran on a CPU.

**Figure 3:**
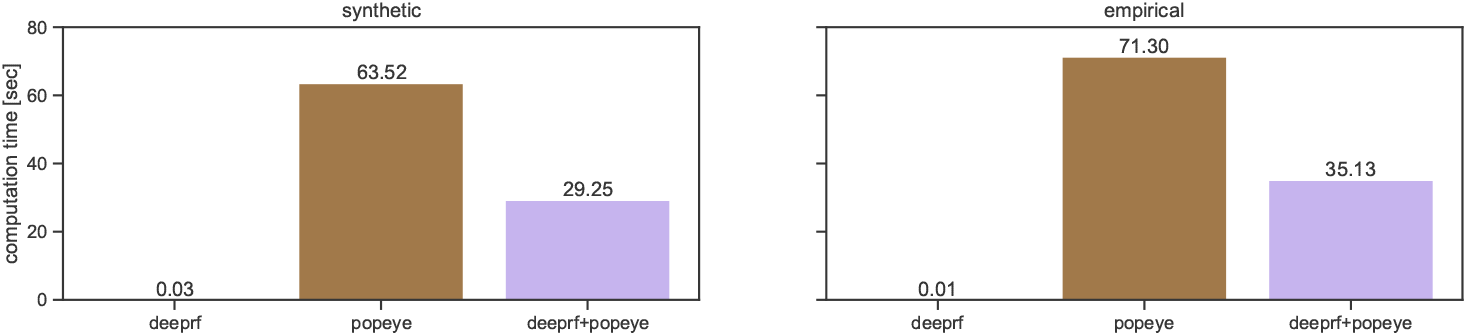
The average computation time in seconds needed per voxel for DeepRF, Popeye and using DeepRF as the coarse fit for Popeye. Note, all models were ran on a CPU.

### 3.2 Validation on simulated data

The parameters as predicted by DeepRF, Popeye and their combination are very close to the target values (see Figure 4). The Pearson correlations of predicted parameters with target values for DeepRF were 0.985, 0.987, 0.991 and 0.986 for *δ*, *σ*, *x*_0_ and *y*_0_ respectively. The same correlations for Popeye were 0.991, 0.988, 0.991 and 0.982 and for the combination of DeepRF and Popeye they were 0.953, 0.975, 0.971 and 0.960. From these values we can observe that all models captured the target values almost perfectly.

**Figure 4:**
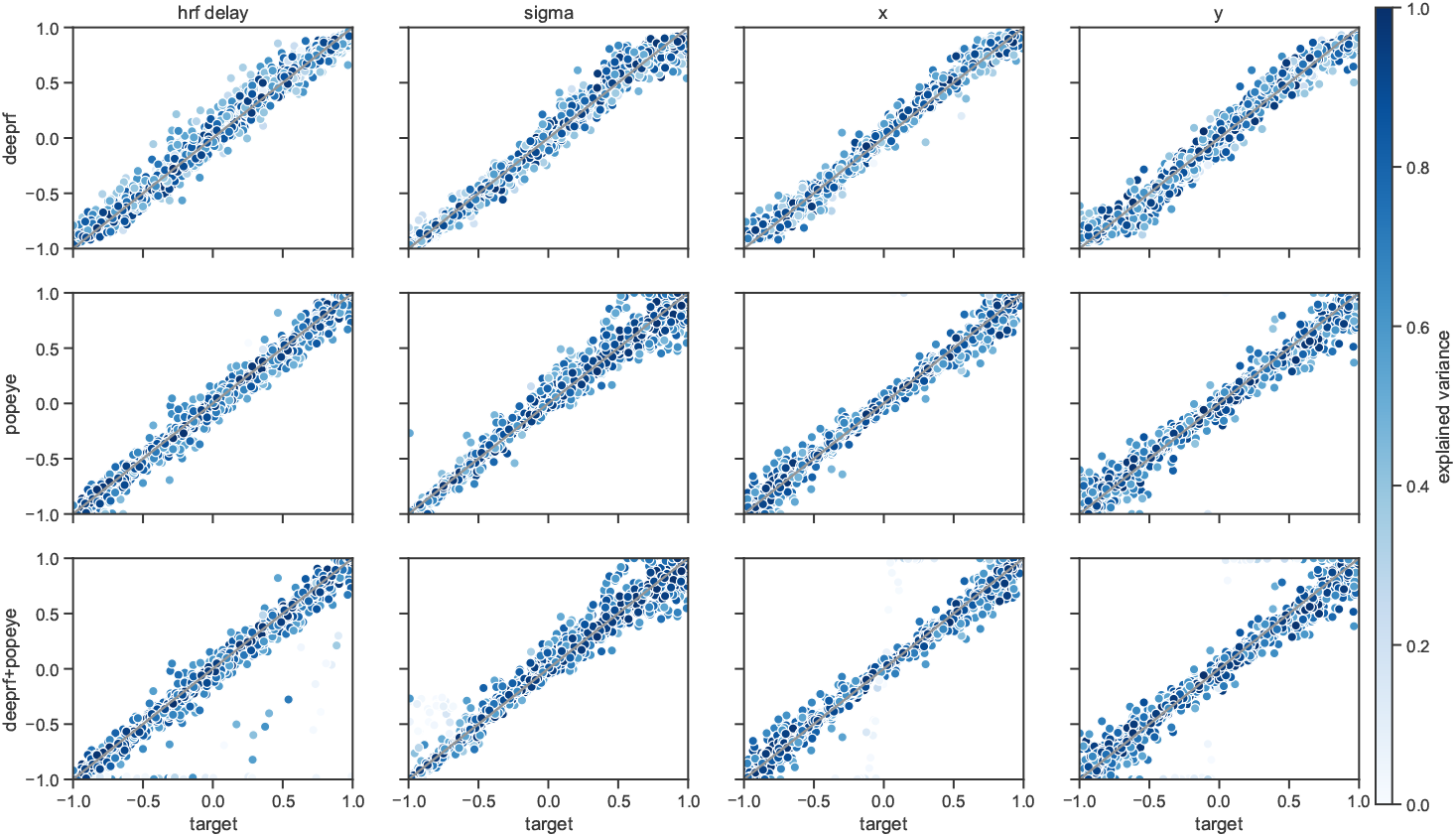
Predicted parameters versus target parameters of the synthetic dataset. The top row shows the predicted parameters of DeepRF versus target, the middle row shows the predicted parameters of Popeye versus target and the last row shows the predicted parameters of DeepRF combined with Popeye versus target. The gray lines indicate the values as they should be given a perfect model. The datapoints represent individual voxels colored by their explained variance.

Given the parameters, one can generate the modeled responses (see Figure 5). We can then compute the explained variance as the squared Pearson correlation between the predicted responses and the observed responses (see Figure 6). The median explained variance was 0.79 for DeepRF, 0.80 for Popeye and 0.80 for their combination. As a comparison, we also computed the median explained variance of the target values, which was 0.80, which denotes the upper bound on the explained variance.

**Figure 5:**
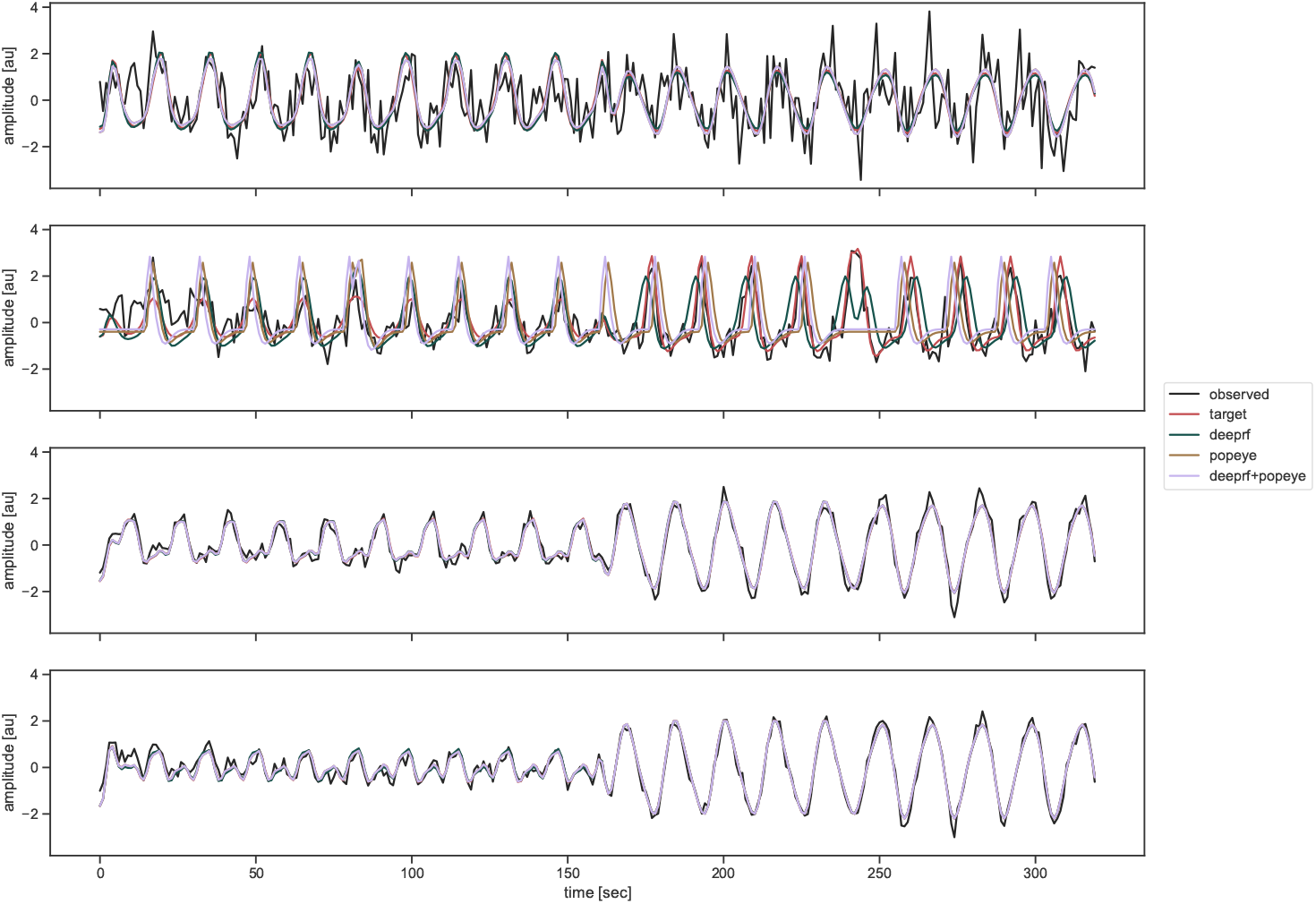
Observed and predicted responses of the synthetic dataset. Four examples of responses are shown where the top two have low explained variance and the bottom two have high explained variance. The black line shows the observed responses which act as input to the prediction models. The target response is the underlying modeled response of the observed data (i.e., the observed data without noise).

**Figure 6:**
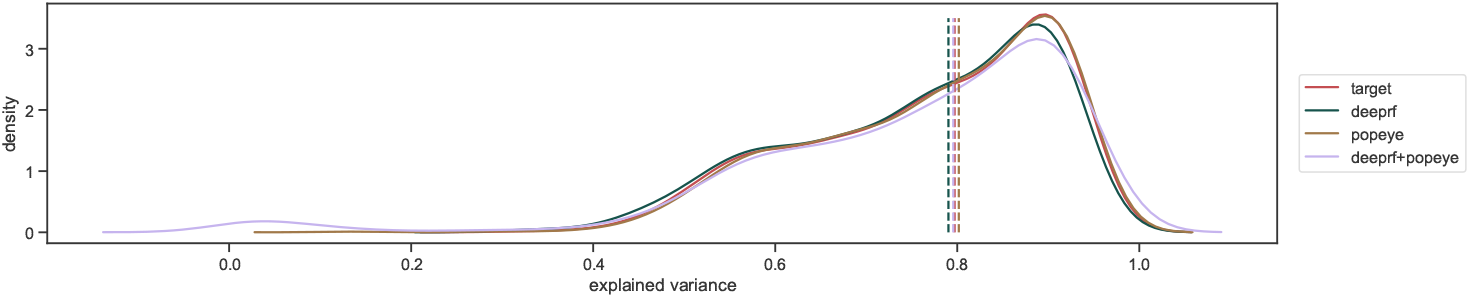
Histograms of explained variances of the synthetic dataset. Solid lines show the distribution of explained variances for the target (i.e., showing the upper limit), DeepRF, Popeye and the combination of DeepRF and Popeye. The dashed vertical lines mark the median values of these distibutions.

To measure the overlap between DeepRF and Popeye we measured the error of both models with the target values as the absolute difference between the predicted parameters and the target parameters, as well as for the difference in explained variance. These were subjected to a paired t-test to test for differences in the error distributions. We found significant effects for the *δ*, *x*_0_ parameters and explained variances (*p* < 0.001), where the the error was always on average larger for DeepRF than for Popeye. The differences were 0.02, 0.00, 0.01 and 0.01 for *δ*, *σ*, *x*_0_ and *y*_0_ respectively and 0.01 for the explained variance. Note that these differences are very small, specifically only 1% of explained variance, which is also reflected in the almost equal response predictions as shown in Figure 5.

### 3.3 Empirical data

As an empirical validation, we applied the models to the Study Forrest dataset. For this dataset, of course, we do not know the ground truth labels. Instead, we take the conventional model, Popeye, as the target model. The parameters as predicted by DeepRF and the combination of Deeprf and Popeye, are very close to the target values of Popeye (see Figure 7). The Pearson correlations of predicted parameters with target values for DeepRF were 0.889, 0.721, 0.975 and 0.985 for *δ*, *σ*, *x*_0_ and *y*_0_ respectively. The same correlations for the combination of DeepRF and Popeye were 0.779, 0.916, 0.972 and 0.988. From these values we can observe that there is a close correspondence between DeepRF and Popeye on empirical data.

**Figure 7:**
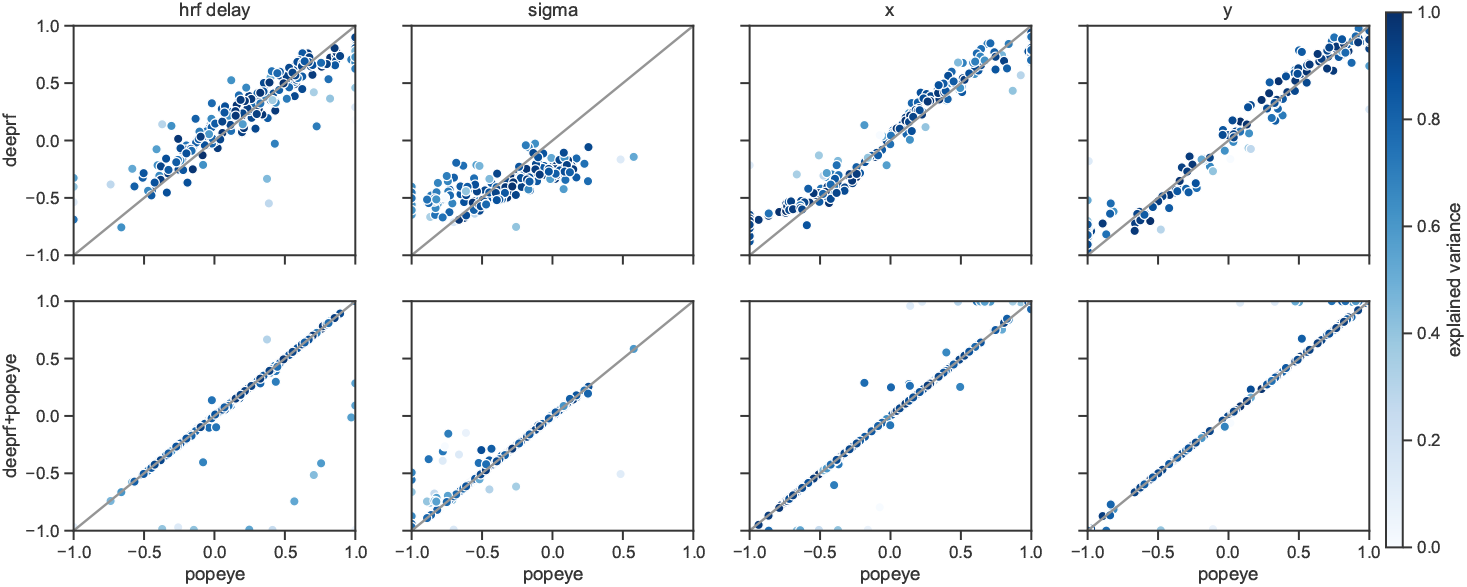
Predicted parameters of DeepRF and DeepRF combined with Popeye versus Popeye parameters on the empirical dataset. The top row shows the predicted parameters of DeepRF, while the bottom row shows the predicted parameters of DeepRF combined with Popeye. The gray lines indicate the values when a model predicts exactly like Popeye. The datapoints represent individual voxels colored by their explained variance.

Given the parameters, one can generate the modeled responses (see Figure 8). We can then compute the explained variance as the squared Pearson correlation between the predicted responses and the observed responses (see Figure 9). The median explained variance was 0.72 for DeepRF, 0.77 for Popeye and 0.74 for their combination.

**Figure 8:**
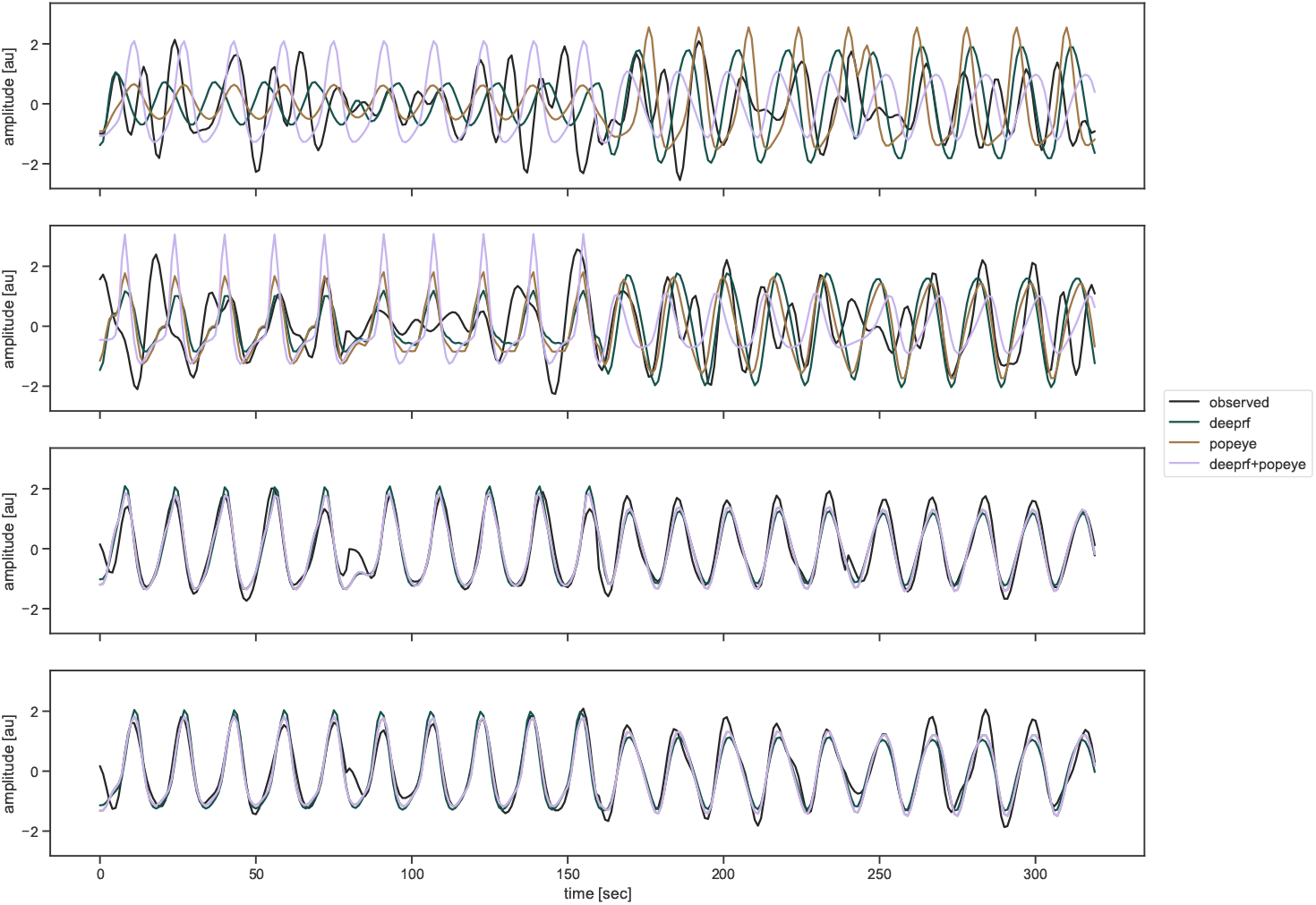
Observed and predicted responses on the empirical dataset. Four examples of responses are shown where the top two have low explained variance and the bottom two have high explained variance. The black line shows the observed responses which act as input to the DeepRF, Popeye and combined models.

**Figure 9:**
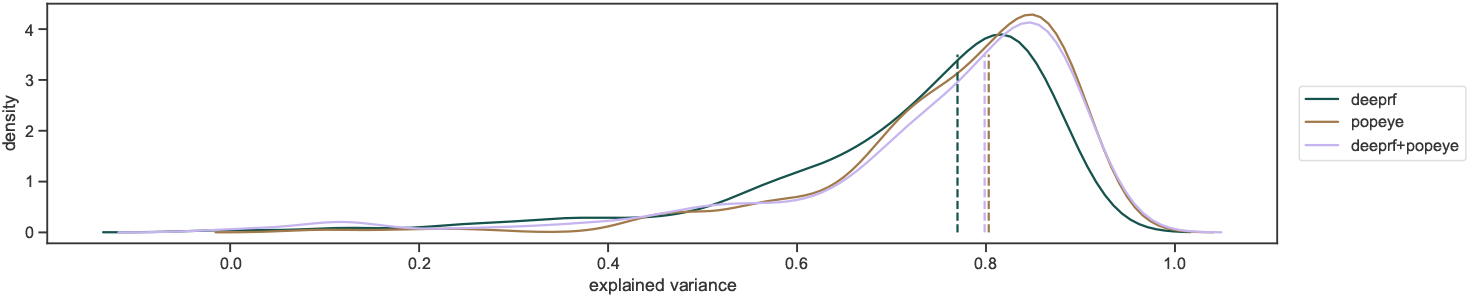
Histograms of explained variances on the empirical dataset. Solid lines show the distribution of explained variances for DeepRF, Popeye and the combination. The dashed vertical lines mark the median values of these distributions.

## 4 Discussion

We introduced DeepRF as a new approach to pRF mapping. Compared to conventional pRF mapping, DeepRF is more than three orders of magnitude faster with only 1% performance loss on average on simulated data and about 5% reduced performance on average on empirical data.

In practice, this entails that we can obtain whole-brain receptive field maps in seconds rather than hours. Note also, that despite the reduced overlap between the models’ *δ* and *σ* parameters, the *x*_0_ and *y*_0_ parameters are very well captured. Hence, DeepRF allows retinotopy, the delineation of visual ROIs (e.g., V1, V2, V3) to be done within seconds, instead of hours. Additionally, our results also imply that pRF models with many more free parameters that could previously not be fitted in a realistic time frame come within reach.

We not only massively improve time efficiency but also greaty reduce the number of computational cycles needed to estimate pRF models, thereby reducing the carbon footprint of these resource-hungry analysis approaches [22, 26]. Note further that the underlying theoretical motivation of our work [1] is fully general. This suggests that a wide range of other demanding analysis techniques within and beyond neuroscience can likewise be improved both in terms of speed and energy demands.

We have to note here, that we performed model comparison on the level of explained variances. One has to be cautious here, as a model that captures noise will gain higher explained variance. Popeye has the power to fit any response well by optimizing its explained variance directly, whereas DeepRF might fail doing so if the signal to noise ratio drops largely or is different between training and testing. In Figure 5 and Figure 8 we show two voxels with high explained variance and two with low explained variance for all models. Note, that the ones with low explained variance typically have higher noise levels.

DeepRF can further be extended to predict posterior distributions over parameters, affording a Bayesian approach to pRF mapping [21]. For example, this can be achieved by adding dropout [12] to the model and using test-time dropout to predict a distribution over parameter values [14]. This may then also capture the noise-related fits discussed above as reduced certainty.

DeepRF might be further improved by allowing the model fitting to be done on predicted responses, as is done in the conventional approach. Currently, DeepRF minimizes the mean squared error between the predicted and target parameters. The loss function might be more informed if with the predicted parameters the error is computed at the level of differences between predicted and observed responses, directly optimizing the explained variance.

Concluding, we have shown that receptive field mapping using DeepRF can be achieved at a speed that is more than three orders of magnitude above the conventional pRF mapping approach. This opens up new possibilities for furthering our understanding of the functional properties of the human brain.

## Acknowledgements

This work has been partially supported by a VIDI grant (639.072.513) from the Netherlands Organi-zation for Scientific Research.

